# Dissecting Oxidative Stress and Organismic Response to various Temperature Regimes in the midge C. riparius

**DOI:** 10.1101/2025.05.28.656541

**Authors:** Burak Bulut, Maximilian Geiss, Markus Bernard, Halina Binde Doria, Barbara Feldmeyer, Markus Pfenninger

## Abstract

Oxidative stress, driven by reactive oxygen species (ROS), poses a major challenge for organisms facing temperature fluctuations. This study provides the first direct *in vivo* measurements of ROS production in an insect, *Chironomus riparius*, across a broad range of ecologically relevant temperatures. We observed a U-shaped pattern of oxidative stress, with minimal ROS levels within an optimal thermal window (12-18°C) and significantly elevated stress at both cold and warm extremes. Crucially, our findings reveal distinct underlying molecular mechanisms for ROS generation at these extremes: at low temperatures, ROS production is predominantly of the superoxide group, linked to hypoxia-induced hemoglobin autoxidation. Conversely, at high temperatures, the hydrogen peroxide group dominates, associated with increased metabolic rate and heat stress signaling pathways. Transcriptomic analysis shows that *C. riparius*’s antioxidant defense system adapts accordingly, selectively upregulating mechanisms to counteract the specific dominant ROS type at different temperatures. This mechanistically differentiated oxidative stress and the modulated organismic response profoundly impacts the overall ecological success and evolution of *C. riparius* as a model for thermal stress in ectotherms.

**Summary Statement:** This study shows that cold and heat stress activate different oxidative damage pathways in midge larvae, explaining how organisms face unique physiological limits in thermal extremes.

## Introduction

In their natural habitats, most organisms are subjected to a range of temperatures throughout their lifespan. Temperature can vary across multiple temporal and spatial scales, ranging from gradual changes across seasons overlain by multi day weather periods and rapid fluctuations over the course of a single day as well as spatial heterogeneity within the habitat. These fluctuations are especially pronounced in temperate regions, where seasonal shifts in temperature can be substantial, requiring organisms to cope with prolonged periods of suboptimal or stressful conditions (Heise et al., 2007). Given that temperature influences the rate of all chemical reactions, each temperature represents a distinct environmental condition that ultimately affects organismal fitness (Huey and Kingsolver, 2011). Deviations from an optimal temperature range the organism is adapted to may therefore result in increased stress, diminished fitness, and, in the extreme, mortality (Gasparrini et al., 2015).

One consequence of temperature-related stress is the production of reactive oxygen species (ROS) (Suzuki and Mittler, 2006) (Figure 1). ROS are highly reactive molecules that necessarily occur in every living cell. The majority of ROS generated within cells are produced by mitochondria as an unavoidable byproduct of the electron transport chain, primarily by complexes I and III. The actual ROS composition produced depends strongly on the stress-inducing conditions and the organism’s condition and may thus vary widely (Das and Roychoudhury, 2014; Guo et al., 2023).

**Figure 1:**
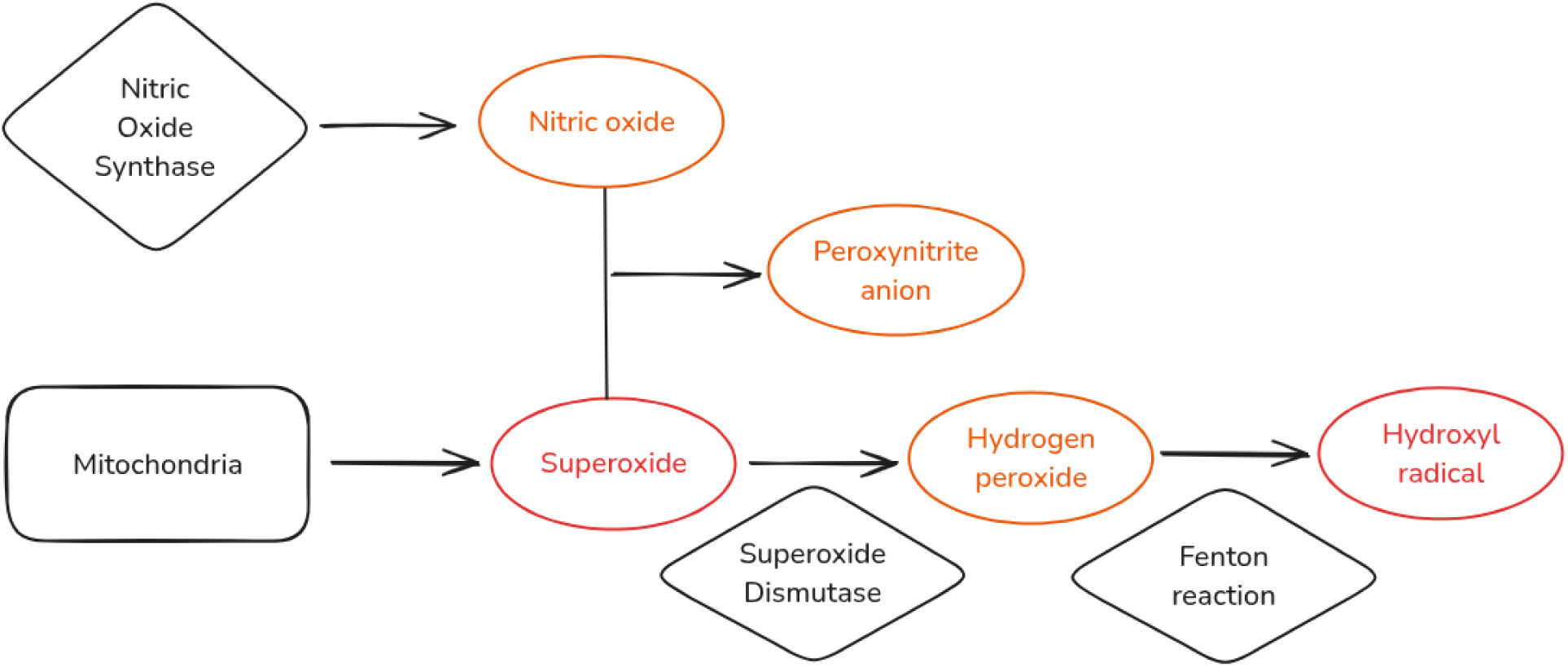
Illustration of the mitochondrial origin and interconversion pathways of reactive oxygen species (ROS) in animals, highlighting specific ROS detected by CellROX Red and CellROX Orange. In animals, the primary ROS produced is superoxide from mitochondria, which is then converted to a range of different ROS. While a part is converted to hydrogen peroxide by superoxide dismutase, others may be transformed to peroxynitrite in the presence of nitric oxide radicals which are mainly produced by nitric oxide synthase family enzyme which can react with superoxide to form peroxynitrite anion (Halliwell et al., 1999). Additionally, hydrogen peroxide may be converted in an additional step to a hydroxyl radical through the Fenton reaction by the endoplasmic reticulum (Liu et al., 2004; Sun et al., 2018; Temple et al., 2005). Red-labeled species indicate oxidants specifically detectable by CellROXRed (e.g., superoxide and hydroxyl radical), while orange-labeled species indicate those uniquely detectable by CellROXOrange (e.g., hydrogen peroxide, peroxynitrite, and nitric oxide).

If ROS are not scavenged, they inevitably cause damage to proteins, lipids, and DNA molecules through unspecific oxidation, which leads to a deterioration of tissues, a reduction in survival, and an impairment of reproduction, which are critical factors for overall fitness (Dowling and Simmons, 2009; Guachalla and Rudolph, 2010). However, certain external or internal circumstances can disrupt this equilibrium, resulting in an increase in cellular ROS levels termed oxidative stress (Davidson and Schiestl, 2001; Larkindale and Knight, 2002; Vacca et al., 2004). To prevent the damage caused by ROS, cells have evolved a sophisticated antioxidant defense system that includes both non-enzymatic and enzymatic components. The non-enzymatic antioxidant system, whose basic concept is to produce bait substances that are oxidised by ROS instead of essential cell components (e.g vitamin C, glutathione etc.) (Chinta and Andersen, 2008; Kalt et al., 2008; Liu et al., 2018). On the other hand, the enzymatic antioxidant system uses diverse enzymes to directly turn ROS into less harmful substances. This system encompasses superoxide dismutase, catalase, glutathione peroxidase, thioredoxin, peroxiredoxin, and glutathione transferase (Birben et al., 2012).

Antioxidant enzymes are distributed throughout various cellular compartments to counteract highly reactive species wherever they occur (Chaudière and Ferrari-Iliou, 1999).

Conversely, ROS molecules are not exclusively harmful but are essential for a multitude of cellular processes, including signaling and the defense against xenobiotic agents. As a result, cells attempt to maintain a delicate equilibrium between ROS production and removal by the antioxidant system and/or scavenging enzymes (Alfadda and Sallam, 2012; Thannickal and Fanburg, 2000). Ultimately, oxidative stress induced by ROS not only causes cellular damage but also drives evolutionary change by introducing mutations into the germ-line that can influence an organism’s adaptability and survival across generations (Dowling and Simmons, 2009).

Studies of temperature-induced ROS effects have predominantly focused on plant systems, resulting in a substantial body of literature on the mechanisms and impacts of oxidative stress (Devireddy et al., 2021; Hasanuzzaman et al., 2020; Hendrix et al., 2023). The few prior studies in animals examining temperature-related oxidative stress employed indirect methods, such as measuring enzyme activity and/or gene expression of antioxidative genes (Jia et al., 2011; Yang et al., 2010; Zhang et al., 2015). Moreover, previous studies investigating ROS production have predominantly focused on high (Farahani et al., 2020; Zhang et al., 2015) or low (El-Saadi et al., 2023; Lebenzon et al., 2023) temperature conditions and only a few have examined both thermal extremes within the same framework (Cui et al., 2011; Yang et al., 2010). To our knowledge, no study has yet systematically assessed ROS responses across a continuous, environmentally realistic temperature gradient.

Given the impact of climate change on the shifting of average and extreme temperatures across all environmental contexts (Rummukainen, 2012), it is imperative to address the existing research gap by conducting a systematic measurement of ROS in vivo across a wide temperature range, thereby enabling a quantitative and qualitative comparison of the ROS spectrum produced and the organismal reaction in terms of changes in gene expression. We use the non-biting midge *Chironomus riparius*, which inhabits small streams and ponds as larvae, and only the adults are airborne. We exposed larvae to a temperature gradient (4°C to 28°C) to observe their stress level as measured by ROS production.

## Material and Method

### Test Animals and Experimental Design

As a widespread freshwater insect, *Chironomus riparius* inhabits rivers, lakes, and ponds across diverse climatic zones, including Hessen, Germany, where our lab population originates. In this region, temperature fluctuates seasonally and geographically, with an annual mean of approximately 9.74°C (Waldvogel and Pfenninger, 2021). Seasonal averages provide a more nuanced perspective: during spring and summer (May to September), the main reproductive period temperatures typically range from 13°C to 18°C, while occasional summer heatwaves or warmer microhabitats may push temperatures higher. In contrast, winter temperatures (November to February) often drop below 5°C, sometimes approaching 2°C (Oppold et al., 2016; Pfenninger et al., 2023).

L3-stage larvae of *Chironomus riparius* were obtained from a long-term maintained and regularly refreshed laboratory culture established from the Hasselbach, Hessen, Germany population (Foucault et al., 2019). A total of 20 L3-stage larvae were collected from this laboratory culture for each treatment and placed in 24-well plates. The plates were filled with 2.5 ml of the medium described in detail in Foucault et al. (2019).

We used a relatively recent method to measure ROS levels in living organisms (Kang et al., 2013). The use of cell-permeable fluorogenic probes allows for the measurement of ROS-related fluorescence intensity within the cells of living organisms. Two different reagents were used to identify the different ROS under different temperatures, CellROX Orange (Thermo Fisher cat. no. C10443) and CellROX Red reagents (Thermo Fisher cat. no. C10422). Both reagents that detect oxidative stress in cells and are suitable for measuring ROS in the cytoplasm of live cells. In the reduced state, the reagents are non-fluorescent. However, following oxidation by ROS, they exhibit fluorogenic signals at 545/565 nm for CellROX Orange and 640/665 nm excitation/emission wavelengths for CellROX Red. CellROX Orange is capable of detecting five distinct ROS, including hydrogen peroxide, hydroxyl radical, nitric oxide, peroxynitrite anion, and superoxide anion. In contrast, CellROX Red only detects two ROS, namely the hydroxyl radical and superoxide anion, which are also part of CellROX Orange. The difference between the CellROX Red and the CellROX Orange signal thus gives a hint on the relative proportion of the two ROS groups produced.

A preliminary experiment with different concentrations of CellROX helped us to determine the final concentration of the reagents which did not lead to death of the larvae but was still sufficient to observe a fluorescent signal. Tested concentrations were 0.5 µl to 2.5 µl (results not shown). In the initial set of measurements, each larva was stained with 0.75 µl of CellROX Orange, while in the subsequent set, 1.25 µl of CellROX Red was used for each larva. Single L3 larvae were placed in a single well of a 24 well-plate (Greiner Bio-One 662160) in a climate chamber with a 16:8 light/dark cycle and a light intensity of 550 lux, without aeration, at the treatment temperature for 24 hours. The study employed seven distinct consecutive temperature regimes, encompassing 4°C, 8°C, 12°C, 16°C, 20°C, 24°C, and 28°C. In total we measured 20 larvae per temperature treatment. Larvae were randomly selected from a continuously breeding population pool, and due to the asynchronous nature of the population, individuals likely originated from different females. Consequently, sampling on consecutive days is not expected to introduce systematic variation.

### Reactive Oxygen Species Measurement

Following 24 hours of treatment, the well plates were placed in a styrofoam box for transportation to the microscope to prevent any abrupt temperature change. ROS level was estimated for a total of 20 larvae per temperature and CellROX dye type. Two larvae had to be excluded from further analysis because they died in the process (one from the 20°C treatment group and one from the 28°C treatment group for CellROX Orange). ROS levels were quantified in live larvae using a ZEISS Axio Imager 2 microscope with 10x magnification. The images were captured using AxioVision Rel. (v. 4.8) with an HXP 120 C fluorescence lamp (Item Number: 423013-9010-000), with maximum light intensity and one setting exposure time. We furthermore employed the "43 HE" (BP 550/25 HE, FT 570 HE, BP 605/70 HE, Item Number 489043-9901-000) filter, which excites blue light at approximately 550 nanometres, transmitting emitted red fluorescence above 570 nanometres and filtering out the remaining blue excitation light, thus registering only red fluorescence around 605 nanometres. As the L3 larva were too large to accommodate imaging of the complete body, the imaging was conducted on the first abdominal segment.

The fluorescence field images were analyzed using the ImageJ Fiji software (version 2.15.0), images were converted to 8-bit grayscale from RGB color images to eliminate color differences to be able to solely calculate light intensity. The CellROX Orange and CellROX Red dyes exhibited differential brightness under identical fluorescence illumination conditions. Consequently, distinct thresholds were applied to the respective reagent treatments. Because the maximum fluorescence light intensity was applied, consequently the background noise (background light) was elevated. Therefore the auto threshold was set too high in order to lower the background noise. The mean values were taken as a measure of fluorescence intensity. It should be noted that the fluorescence intensity measured does not necessarily represent the actual amount of ROS present within the cell. Rather, it reflects the current amount of reagent that has entered the cell and undergone oxidation by binding to ROS. The data were subsequently subjected to analysis in R.

### Gene Expression Analysis

A total of 45 L3 larvae were collected from the lab culture for each temperature. The larvae were exposed to the above mentioned temperature treatments in glass bowls filled with sand and medium, as previously described (Foucault et al., 2019). The larvae were provided with aeration and were permitted to feed at their discretion. As developmental time depends on temperature in this ectothermic organism, the duration of exposure varies according to temperature: 25°C for 2 days, 20°C for 3 days, 15°C and 10°C for 4 days, and 5°C for 5 days. For each temperature treatment, multiple replicates of larvae were utilized, with 12 individuals per 25°C, 15°C, and 10°C groups, and 11 individuals per 20°C and 5°C groups. For each individual larva, a separate RNA library was prepared. RNA was extracted using the Direct-zol RNA Miniprep Kits (R2053, Zymo Research) according to the manufacturer’s manual. Library preparation and sequencing was conducted at Novogene on a NovaSeq X Plus instrument.

### RNA Data Analysis

The FASTQ files were aligned to the *C. riparius* reference genome v4 (Pettrich et al., 2024) using HISAT2 (v. 2.1.0) (Kim et al., 2019). Subsequently, the SAM files were sorted and converted to BAM format using Samtools (v. 1.20) (Li et al., 2009). The read counts were extracted per individual using FeatureCounts (version 2.0.6) (Liao et al., 2014).

We subsequently normalized the raw counts using DESeq2 (v. 3.19) (Love et al., 2014) Then excluded genes with read counts <10 in at least 4/5 of all tested temperatures. This resulted in the selection of 118 genes. In this study, we focused on ROS-related genes and their specific reaction norms to different temperatures only. To this end, protein sequences of 134 ROS-related proteins from the model organism *Drosophila melanogaster* were downloaded from UniProt on 30 June 2024. Subsequently, the *C. riparius* genome was queried against the downloaded protein database using BLASTx (v. 2.12.0) (Camacho et al., 2009) to identify and extract the corresponding genes. With this approach, nine previously not annotated genes were discovered in the *C. riparius* genome and incorporated into the analysis. The *C. riparius* protein set was queried against the UniProtKB protein database using the InterProScan (v. 5.52) software to obtain the gene name.

### PCA Analysis and Visualization

To explore coordinated temperature-dependent responses in the antioxidant system, we conducted a principal component analysis (PCA) exclusively on 23 genes known to be directly involved in ROS production or scavenging (Ryan et al., 2020; Zou et al., 2000). The goal was to identify dominant patterns of variation in gene expression across the temperature gradient and to uncover how antioxidant systems respond collectively under thermal stress. The normalized gene expression data for these antioxidant genes were imported into R and pre-processed by applying z-score normalization. PCA was performed using the ‘prcomp()’ function to capture the major axes of variation in antioxidant gene expression across temperature treatments. Sample metadata, including temperature, were extracted from sample identifiers and merged with PCA scores, while gene loadings were obtained from the PCA rotation matrix and combined with ROS target annotations.

Additionally, a broken stick model was used to assess the significance of the variance explained by each principal component.

Visualization of the PCA results was carried out using the ggplot2 package (Wickham, 2011). A biplot was generated, displaying samples in the PC1-PC2 space with points colored by temperature and gene loadings represented as scaled arrows. Complementary boxplots were also created to compare the distribution of principal component scores across temperature treatments, with custom color scales applied for clarity.

All additional statistical analyses, including principal component analysis, broken stick modeling and data visualization were performed in R version 4.4.1 (R Core Team, 2024).

## Results

### Temperature Dependent Stress Levels Across Temperatures Detected via ROS

The stress levels of larvae as measured by the relative fluorescence intensity (RFI) of ROS exhibited a significant temperature dependence (ANOVA p-value < 2.2e-16), and was highest at the two temperature extremes and lowest between 12°C and 18°C which indicates an optimal temperature range (Figure 2). Fitting a second-order polynomial regression model to the data revealed a U-shaped relationship between temperature and RFI (goodness of fit: R^2^=0.6, p-value < 2.2e-16; formula: 44.91672 + 30.85863 * x + 298.3746 * x^2). The polynomial regression model fitted better to the data compared to the linear model according to AIC values of 1250 and 1377 respectively.

**Figure 2:**
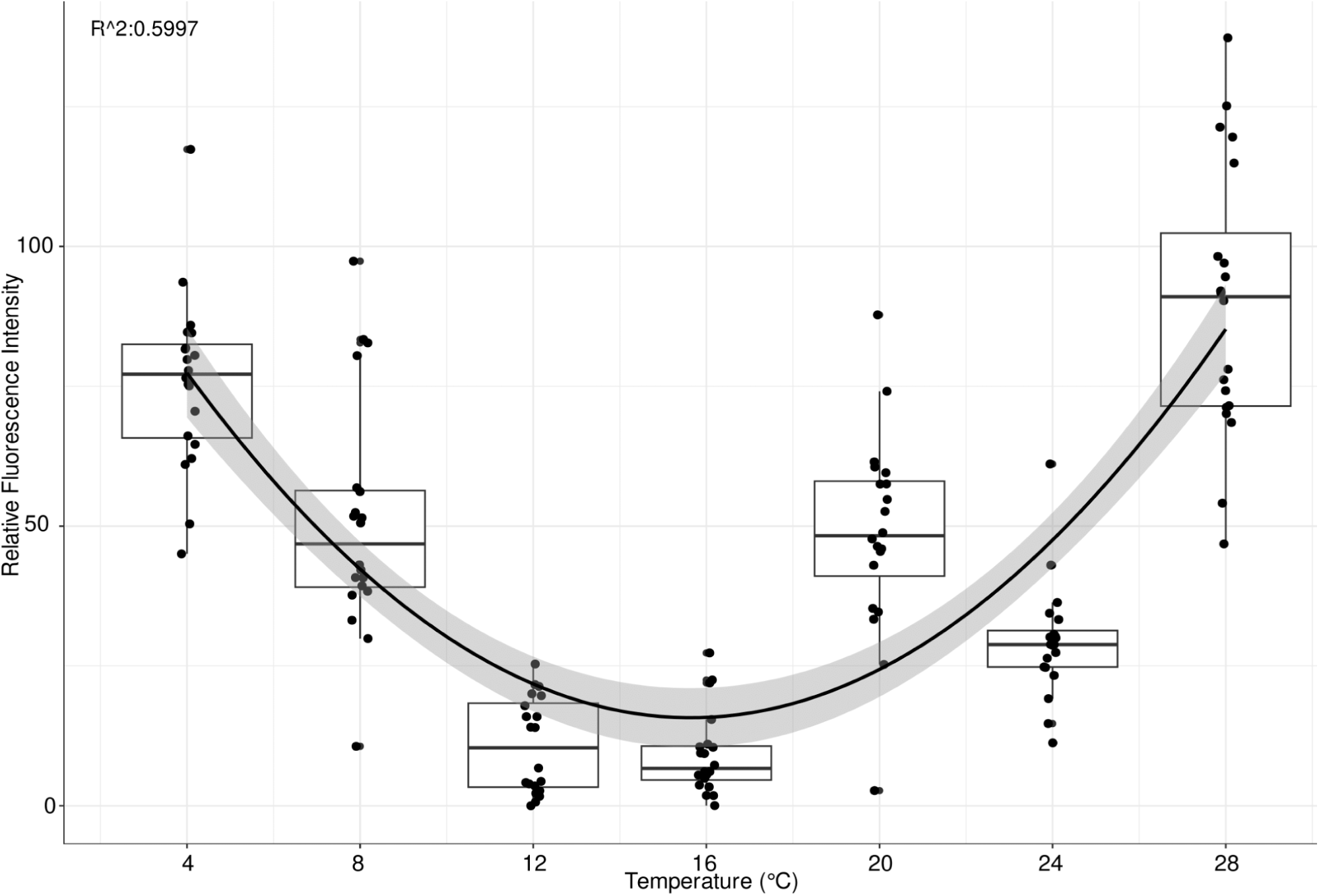
Boxplot depicting the relative fluorescence intensity (RFI) across different temperatures based on CellROX Orange measurements. The distribution of RFI across temperatures fits a parabolic distribution (R^2^ = 0.5997; F-statistic: 105.1 on 2 and 137 DF, p-value: < 2.2e-16) The line shows the fitted regression with a 95% confidence interval (shaded area).

### Characterization of ROS Types Under Different Temperature Conditions

When looking at the relative abundance of ROS species across temperatures, it becomes evident that the abundance of the different ROS-groups, i.e. the ratio of the superoxide group to the hydrogen peroxide group changes with temperature. At 4°C, the superoxide group (hydroxyl radical and superoxide anion; CellROX Red) predominated. In contrast, at 24°C and 28°C, there is an elevated level of the hydrogen peroxide group (hydrogen peroxide, nitric oxide, and peroxynitrite; CellROX Orange-Red). At optimal temperatures, the amount of ROS is lower than at the extremes and is dominated by the superoxide group (Figure 3).

**Figure 3:**
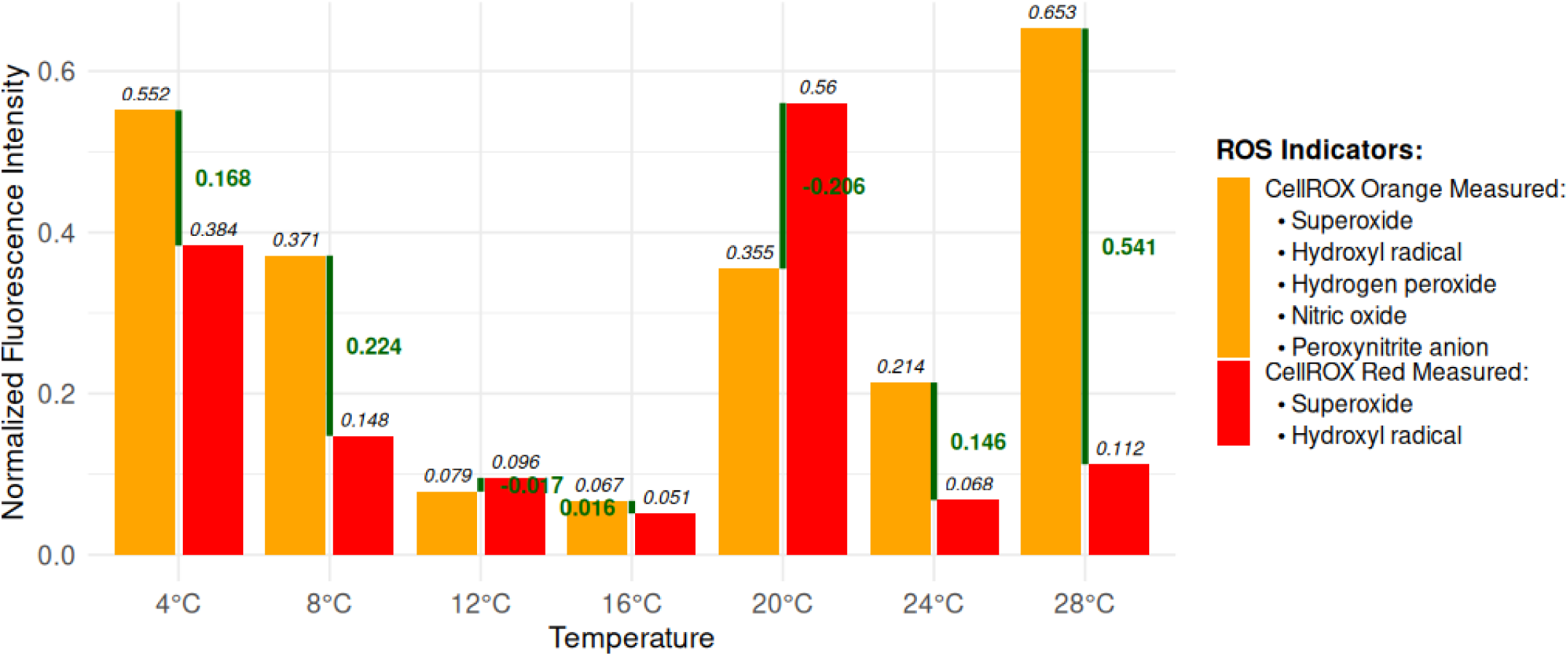
Normalized fluorescence intensity of ROS indicated by orange bars (total measured ROS by CellROX Orange) and red bars (subset of ROS by CellROX Red) at temperatures ranging from 4°C to 28°C. Total measured ROS include superoxide, hydroxyl radical, hydrogen peroxide, nitric oxide, and peroxynitrite anion. The subset of ROS consists of superoxide and hydroxyl radicals. All values are normalized. Green segments, with corresponding numerical values, illustrate the difference between total and subset ROS levels (CellROX Orange - CellROX Red), representing the combined contribution of hydrogen peroxide, nitric oxide, and peroxynitrite anion.

Under optimal conditions, only a low level of ROS detected by CellROX Red is produced, while the additional oxidant species detected by CellROX Orange become prominent under stress. Notably, when 16°C data was used as a baseline, the normalization method proved inconsistent, as seen at 12°C and 20°C where the Red signals unexpectedly exceeded the Orange, pointing to a methodological rather than strictly biological discrepancy. This suggests that our baseline subtraction approach, while useful under some conditions, does not adequately account for the overlapping sensitivities of the probes.

### Principal Component Analysis of ROS Associated Genes Under Different Temperature Conditions

Principal component analysis (PCA) was performed on normalized read counts of 23 antioxidant genes directly related to either ROS production or scavenging to determine temperature-dependent expression patterns. Significant PCs were revealed using the Broken Stick model (Sup. Data 1), where the first 3 principal components (PC) capture different aspects of gene expression variation, with PC1, PC2, and PC3 explaining 24.2%, 16.2%, and 10.8%, respectively. Together, they explained more than 50% of the data (Figure 4 A).

**Figure 4:**
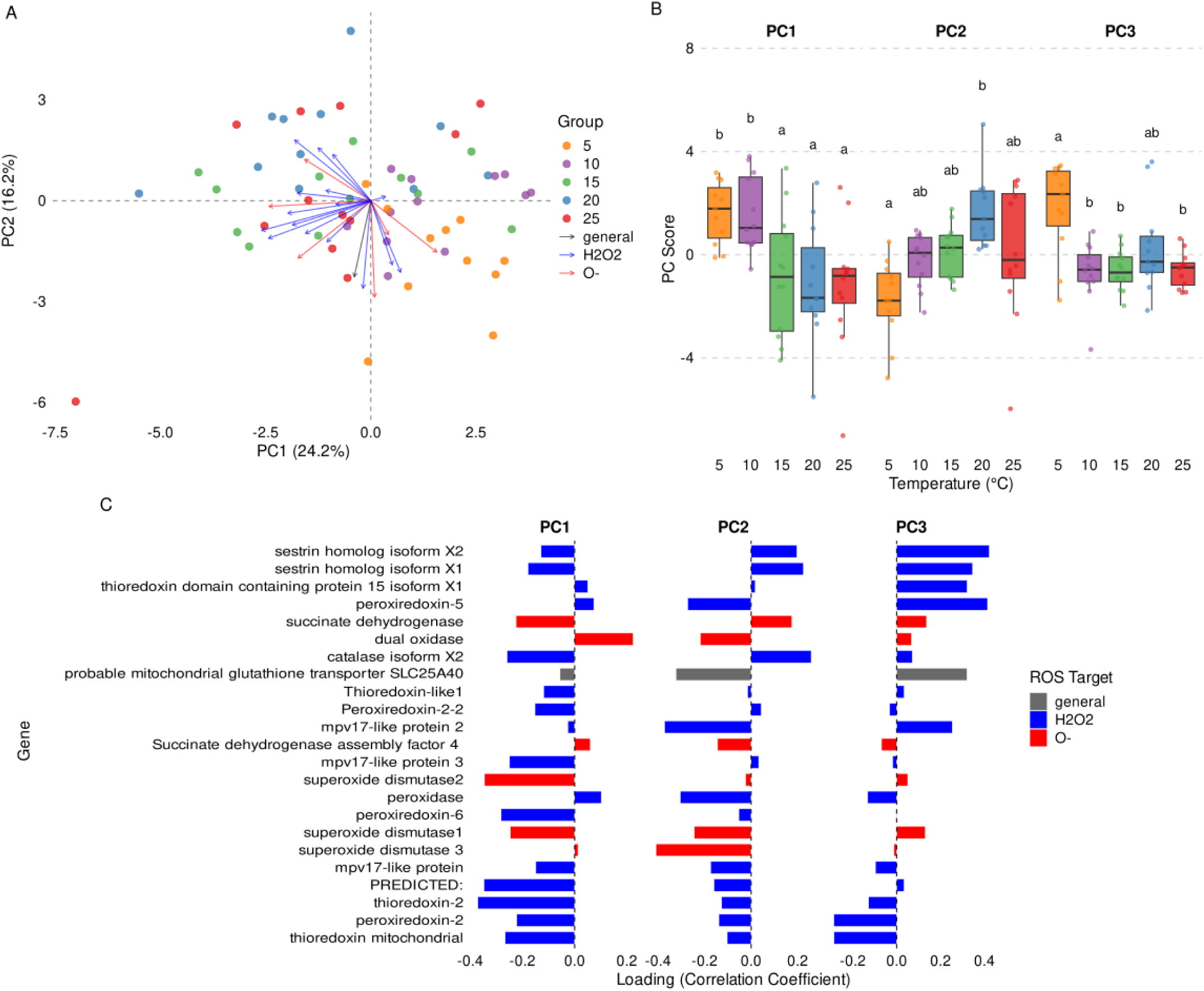
PCA of antioxidant gene expression across five temperatures, showing temperature-specific patterns and the contribution of ROS-targeted genes to variation along principal components. (A) Biplot of the first two principal components derived from a principal component analysis (PCA) of 23 antioxidant gene expression profiles across five temperature treatments (5°C, 10°C, 15°C, 20°C, 25°C). Samples are color-coded by temperature, and vectors indicate each gene’s contribution to the observed variation along PC1 and PC2. Genes are labeled and colored according to their primary ROS target, with gray for general (non-specific) antioxidants, blue for hydrogen peroxide-related antioxidants, and red for superoxide-related antioxidants. (B) Boxplots of sample scores for PC1, PC2, and PC3 across all temperature treatments, illustrating shifts in overall gene expression patterns. Letters indicate statistically significant differences among temperatures within each principal component (Tukey’s HSD test, p < 0.05). (C) Contribution of 23 ROS production or scavenging candidate genes to PC1, PC2, and PC3. Genes are again categorized by primary ROS target, using the same color scheme as in panel A.

The PCA biplot showed a clear temperature-dependent clustering of the samples. The low-temperature samples (5-10°C) formed distinct clusters in the positive PC1 region, while the high-temperature samples (20-25°C) clustered in the negative PC1 region. Intermediate temperature samples (15°C) occupied transitional positions between these clusters, indicating a gradual shift in gene expression patterns across the temperature gradient.

Analysis of PC scores across temperatures revealed systematic variation in gene expression patterns. PC1 scores showed a strong correlation with temperature, transitioning from positive values at low temperatures to negative values at high temperatures. PC2 scores showed another clear distinction between low temperature (5°C) and high temperature (20-25°C), while the 10 and 15°C data sets fell in the middle. This suggests a threshold of regulation in the antioxidant response system. PC3 provided additional information on cold-temperature-specific responses, particularly in the 5°C data (Figure 4 B) (Sup. Data 1).

### Gene Contributions to Temperature Response

Based on the PCA (Figure 4 A) we obtained gene loadings to determine the contribution of the 23 antioxidant genes directly related to either ROS production or scavenging to the temperature response. In PC1, thioredoxin-2, two mitochondrial thioredoxin isoforms, two superoxide dismutase variants, peroxiredoxin-6, catalase isoform X2, and mpv17-like protein 3 showed strong negative loadings (–0.2 to –0.4), indicating higher expression levels at elevated temperatures. In contrast, dual oxidase exhibited a positive loading, suggesting increased expression at lower temperatures.

PC2 and PC3 captured variation associated with cold responses. Strong negative loadings in PC2 were observed for superoxide dismutases, mpv17-like protein 2, mitochondrial glutathione peroxidase, and peroxiredoxin-5, indicating their greater contribution under cold conditions. Positive PC2 loadings for catalase isoform X2 and sestrin homologs suggested differential antioxidant activation at intermediate temperatures. PC3 was dominated by high positive loadings from sestrin isoforms, peroxiredoxin-5, and thioredoxin domain-containing protein 15, indicating a specific role for hydrogen peroxide detoxification at low temperatures (Figure 4 C) (Sup. Data 1).

## Discussion

Organisms need to be able to cope with a variety of different temperatures across different time scales and magnitudes. Even though water bodies may buffer temperature fluctuations to certain extent, water bodies in temperate regions, and especially small water bodies change temperatures on a daily and definitely seasonal time scale. Thus organisms inhabiting these regions need to be adapted to cope with conditions, but they are still vulnerable to temperature extremes. Temperature extremes exert strong stress on organisms, however the detection and quantification of this stress response is not always easy. This study presents the first direct *in vivo* measurements of ROS under a broad range of temperatures in an insect. In contrast to earlier studies that employed indirect methods, such as enzyme activity assays or in vitro ROS measurements (Cui et al., 2011; Jia et al., 2011; Yang et al., 2010; Zhang et al., 2015), our direct in vivo measurements of ROS provide a more comprehensive and precise understanding of ROS production in response to a range of temperatures encountered by the species in their natural habitat.

### Temperature-Dependent Stress Response

Our findings demonstrate that oxidative stress in *C. riparius* follows a U-shaped pattern across the temperature gradient, with elevated ROS levels at both upper and lower temperature extremes. The increased ROS levels at the extreme temperatures align with observations in other ectotherms, where temperature extremes challenge cellular homeostasis. For instance, in ectothermic species like fish and amphibians, either heat or cold stress can lead to increased ROS (Paital and Chainy, 2014). This phenomenon can be explained by the interplay of several physiological factors, including temperature-dependent changes in metabolic rate, mitochondrial activity, and gene expression patterns (Bury et al., 2018; Vinagre et al., 2014).

At high temperatures, the metabolic rate increases, leading to higher oxygen consumption and elevated ROS production as a metabolic by-product (Pamenter et al., 2018; Rollins-Smith and Le Sage, 2023). Consistent with the generally increased metabolic rate observed at higher temperatures (Sahragard and Rafatifard, n.d.; Schulte, 2015), we identified linearly increasing reaction norm of five NADH dehydrogenase genes and exponential increase of a cytochrome c reductase gene with increasing temperature indicating enhanced expression of components within complexes I and III of the electron transport chain. This upregulation likely reflects a response to elevated energy demands. As temperatures rise, cellular ATP requirements increase, driving the need for efficient mitochondrial ATP production. The observed increase in mitochondrial gene transcription may therefore serve to increase the flow of electrons through the electron transport chain to meet these heightened energy demands under thermal stress (Park and Kwak, 2014).

On the other hand, at lower temperatures, mitochondrial function is disrupted as enzyme function slows down and oxygen consumption is decreased, leading to increased electron leakage from the mitochondrial electron transport chain and subsequent enhancement of ROS production (Jørgensen et al., 2023). In accordance with this, we observe reduced expression of complex I and III genes at lower temperatures, which may represent a compensatory metabolic adjustment rather than dysfunction, aligning with diminished energy requirements in cold conditions. Downregulation of these genes in cold conditions is likely a strategy to conserve energy and minimize ROS production by reducing the electron transport chain’s activity, which is particularly useful in low-temperature environments where ATP demand decreases (Colinet et al., 2017).

### Different ROS Profiles Across the Temperature Gradient

Beyond the observed U-shaped stress response, our approach of using different CellROX reagents further disentangle these responses: at low temperatures, the superoxide group (superoxide and hydroxyl radical) predominates, whereas at high temperatures the hydrogen peroxide group (hydrogen peroxide, peroxynitrite anion and nitric oxide) dominates.

At low temperatures (e.g., 4°C), the elevated ROS production is largely driven by hypoxia-induced hemoglobin autoxidation. In *C. riparius* larvae, hemoglobin levels are known to be high and increase further under hypoxic conditions (Weber, 1980). Under such low-oxygen environments, reduced oxygen diffusion promotes autoxidation of oxyhemoglobin to methemoglobin (Lushchak et al., 2005; Lushchak and Bagnyukova, 2007), a process that generates superoxide radicals and, to a lesser extent, hydrogen peroxide (Anand et al., 2013). Furthermore, diminished peroxidase activity under these hypoxic conditions favors the Fenton reaction, thereby forming hydroxyl radicals. Such molecular mechanisms not only explain the ROS profile observed at low temperatures but may also account for the associated developmental delays observed below 15°C (Abele and Puntarulo, 2004; Pfenninger and Foucault, 2020).

At elevated temperatures, the predominance of the hydrogen peroxide group reflects both a heightened metabolic rate and specific heat stress signaling pathways. As temperature increases, the consequent elevated ATP demand drives greater electron leakage in the electron transport chain, leading to increased production of superoxide anions. Upregulated superoxide dismutase activity then converts these anions into hydrogen peroxide (Murphy, 2009). Beyond this general metabolic response, heat stress triggers additional molecular responses such as increased nitric oxide production through the modulation of nitric oxide synthase that promote the formation of peroxynitrite when superoxide reacts with nitric oxide (Kirsch and de Groot, 2002; Müller, 1997).

Ecologically, these molecular processes have profound implications for *C. riparius* populations. Elevated temperatures can accelerate larval development, thereby reducing generation time, which principally increases fitness (Foucault et al., 2019). However, the trade-off is significant: higher temperatures compromise larval survival, reduce pupation success, and thus overall impair adult reproductive fitness. Experiments have shown that while optimal growth and relatively short generation times are maintained between 16°C and 23°C (Oppold et al., 2016), exposure to temperatures around 28°C leads to a notable decline in population growth (Nemec et al., 2013). This suggests that the benefits of accelerated growth rates at elevated temperatures could be counterbalanced among others by the physiological costs of the increased ROS production shown here, ultimately affecting the species’ long-term viability and ecological success.

Our findings likely also have evolutionary implications for the species. The mutation rate response across temperatures reported by Waldvogel and Pfenninger (2021) exhibited an optimality pattern, superficially similar to the one observed in our ROS measurements.

Observed mutation rates had their minimum at ∼18°C and increased with increasing temperatures, but also towards colder temperatures, reaching a maximum at 12°C (Waldvogel and Pfenninger, 2021). Below this temperature, reproduction is not possible anymore in *C. riparius* and therefore, transgenerational mutation rates could not be measured. However, ROS levels had their minimum at 12°C and are therefore likely not the major cause for the increased mutation rate observed at this temperature. This suggests that the extended generation times at cold temperatures (Oppold et al., 2016) allows even low ROS levels more time to inflict DNA damage (Beal et al., 2019) and thus increase mutation rates. However, the overwintering generation of *C. riparius* in nature is exposed for several months to temperatures down to 4°C (Pfenninger et al., 2023; Waldvogel and Pfenninger, 2021). The increased ROS level at these cold temperatures observed here could thus still be a major driver of mutagenesis in natural populations, due to the interplay of increased oxidative cold stress and extended exposure time during hibernation.

### Molecular Mechanisms of Antioxidant Defense Across Temperature Gradients

Our analysis revealed a targeted molecular response by which *C. riparius* reacts to temperature-induced oxidative stress, corroborating the stress level results observed above.

At higher temperatures (20-25°C), we observed elevated expression of hydrogen peroxide-neutralizing systems, including thioredoxins, peroxiredoxin-6, and catalase isoform X2 (Figure 4 C). This shift toward hydrogen peroxide management likely reflects the increased mitochondrial activity and metabolic rate at elevated temperatures, which typically generate greater quantities of this particular ROS (Pamenter et al., 2018; Rollins-Smith and Le Sage, 2023). The concurrent upregulation of superoxide dismutase variants suggests a coordinated, multi-layered defense strategy addressing both primary (superoxide) and secondary (hydrogen peroxide) ROS species.

Conversely, colder conditions (5-10°C) triggered a distinctly different antioxidant profile. While the increased expression of dual oxidase at lower temperatures indicates enhanced hydrogen peroxide production under cold conditions, our RFI data reveal that superoxide anion emerges as the dominant oxidant in this thermal range. This dual oxidase activity, despite contributing to hydrogen peroxide levels, appears secondary to the more significant superoxide challenge. The cold-temperature response therefore appropriately emphasizes superoxide management, with complex regulation of superoxide dismutases alongside hydrogen peroxide scavengers. This seemingly counterintuitive production of multiple ROS species at low temperatures may serve critical signaling functions, potentially activating cold-adaptive pathways or maintaining cellular redox homeostasis despite reduced metabolic activity (Jørgensen et al., 2023; Koštál et al., 2007; Murphy, 2009). The sophisticated balance between different ROS species at metabolic extremes suggests a highly evolved regulatory system that precisely calibrates oxidative defense according to specific thermal challenges.

### Conclusion and Future Directions

This study provides the first direct *in vivo* evidence for an optimality pattern of overall oxidative stress across a broad temperature range in the insect *Chironomus riparius*. Minimal reactive oxygen species (ROS) levels were observed within an optimal thermal window (12-18°C) and significantly elevated stress at both cold and warm extremes. This overall pattern arises from distinct underlying mechanisms: ROS production of the superoxide group (superoxide, hydroxyl radical) predominates at low temperatures, primarily linked to hypoxia-induced hemoglobin autoxidation, while the production of the hydrogen peroxide group (hydrogen peroxide, peroxynitrite, nitric oxide) dominates at high temperatures, associated with increased metabolic rate and heat stress signaling. Notably, while the specific production profiles of these ROS groups do not individually form a U-shape, their combination results in the observed stress minimum at 12-16°C. Correspondingly, the molecular response of the plastic expression of antioxidant genes, is specifically tailored to counter the dominant ROS type prevalent at different temperatures rather than simply following the overall ROS stress. These findings establish that temperature extremes impose significant, but mechanistically different, forms of oxidative stress, impacting larval development, survival, and overall fitness. Our findings suggest that the increased ROS production at both the cold and warm ends of the temperature range experienced in natural habitats strongly influences the ecology and evolution of the species.

## Acknowledgements

Burak Bulut thanks to DFG for support by a grant to MP (PF 390/15-1).

## Data Availability

The data is accessible at the European Nucleotide Archive (ENA) under accession number PRJEB89193.

## CRediT author statement

**Burak Bulut:** Conceptualization, Data curation, Formal analysis, Methodology, Visualization, Writing - original draft; **Maximilian Geiss:** Conceptualization, Data curation, Formal analysis **Markus Bernard:** Methodology **Halina Binde Doria:** Methodology, Conceptualization **Barbara Feldmeyer:** Conceptualization, Data curation, Writing - review & editing; **Markus Pfenninger:** Conceptualization, Methodology, Validation, Resources, Project administration, Writing - review & editing, Funding acquisition.

